# Cardiac Development Long non-coding RNA (CARDEL) is activated during human heart development and contributes to cardiac specification and homeostasis

**DOI:** 10.1101/2023.02.19.529122

**Authors:** Isabela T. Pereira, Rubens Gomes-Júnior, Aruana Hansel-Frose, Man Liu, Hossam A.N. Soliman, Sunny S.K. Chan, Samuel C. Dudley, Michael Kyba, Bruno Dallagiovanna

## Abstract

Successful heart development depends on the careful orchestration of a network of transcription factors and signaling pathways. In recent years, the *in vitro* cardiac differentiation using human pluripotent stem cells (hPSCs) has been used to uncover the intricate gene network regulation involved in the proper formation and function of the human heart. Here, we searched for uncharacterized cardiac developmental genes by combining a temporal evaluation of the human cardiac specification *in vitro* with the analysis of fetal and adult heart tissue gene expression. We discovered that *CARDEL* (CARdiac DEvelopment Long non-coding RNA; LINC00890; SERTM2) expression coincides with the commitment to the cardiac lineage. *CARDEL* knockout hPSCs differentiated poorly in cardiac cells, and hPSC-derived cardiomyocytes showed faster beating rates after *CARDEL* controlled overexpression during differentiation. Altogether, we demonstrate physiological and molecular evidence that *CARDEL* expression contributes to sculpting the cardiac program during cell-fate commitment.

## Introduction

The delivery of oxygen and nutrients to the many tissues throughout the body is possible due to the function of a complex muscular organ, the heart. The function of the heart depends on the interplay of different cell types, including cardiomyocytes, smooth muscle cells, endothelial cells and cardiac fibroblasts (Meilhac and Buckingham, 2018; Spater et al., 2014). Cardiomyocytes are the fundamental work unit of the heart, which ensure contraction of the chambers and efficient blood flow. To make a specialized cardiomyocyte is a complex process. During embryonic development, the commitment to the cardiac lineage includes several steps that are regulated by a network of transcription factors and signaling pathways, which control the specification and maturation of cardiac cells (Meilhac and Buckingham, 2018; Olson, 2006; van Vliet et al., 2012).

The cardiac tissue is mostly derived from the mesoderm layer, which is one of the three distinct germ layers specified from the embryo inner cell mass. Lineage-tracing and molecular profiling experiments have provided evidence that cardiovascular progenitors are specified early in development, mostly at the stage of mesoderm induction (Protze et al., 2019). Animal models have made important contributions to our knowledge of cardiac developmental events, especially those underlying the early genetic pathways (Evans et al., 2010). However, genomic differences among species that directly impact development regulation has become much clearer. Moreover, species differences in heart physiology, such as heart rate, calcium and potassium currents, and also the myosin composition, make modeling human diseases in mice challenging (Feng and Jin, 2018; Sadayappan et al., 2009; Su et al., 2003). The limited suitability of animal models to address human-specific aspects of development, disease and therapy emphasizes the importance of human specific studies. In addition to the restricted access to human fetal-derived biological material, the advantages of using human cardiomyocyte-based platforms for drug discovery, predictive toxicology and cell therapy promoted the rapid implementation of human pluripotent stem cells (hPSC) differentiation protocols (Devalla and Passier, 2018; Hofbauer et al., 2021; Leitolis et al., 2019).

The careful orchestration of several signaling pathways and regulatory genes is crucial for successful heart development and, therefore, deserve special attention as defects in these early cell fate decisions might contribute to congenital heart diseases or, in severe cases, stillbirth (Buijtendijk et al., 2020; van Vliet et al., 2012). One key advantage of *in vitro* cardiac differentiation experiments is to recapitulate key developmental events that become readily accessible to study. Associated with a variety of “omics” assays, it represents a powerful approach to uncover new players in cardiac specification. In this report, we describe the strategy to find and functionally characterize the cardiac developmental gene *CARDEL* (CARdiac DEvelopment Long non-coding RNA). We combined a temporal evaluation of the human cardiac specification *in vitro* with an analysis of fetal and adult heart tissue gene expression to search for uncharacterized genes. *CARDEL* knockout and controlled overexpression in hPSCs were accomplished using two distinct and independent cell lines. These complimentary approaches show that *CARDEL* is required for a successful cardiac specification as well as final cardiomyocyte physiological function, establishing *CARDEL* as a critical cardiac gene.

## Results

### *CARDEL* is expressed during the human cardiac lineage specification

Taking advantage of our previously published RNA-seq dataset (Pereira et al., 2018), we narrowed our search for cardiac developmental genes looking at the differentially expressed genes between mesoderm and cardiac progenitor (CP) stages in the *in vitro* differentiation (IVD) of hPSC. Analysis of publicly available RNA-seq of human fetal and normal adult heart tissues was also performed as a second step to limit the genes to those mainly expressed during heart development (Figure 1A). Genes that showed upregulation in CP cells (1354 genes, Table S1-A) and were found as enriched in fetal heart tissues (3503 genes, Table S1-B) were considered potential candidates (intersection = 151 genes, Table S1-C). Among those genes are well-known fetal-enriched genes, such as TNNI1, which is the fetal isoform expressed in cardiac and skeletal muscle during early development (Bedada et al., 2014). As our primary goal was to find novel and uncharacterized genes, Gene Ontology analysis was used to filter out genes assigned to known biological processes (Table S2). Following these criteria (Figures 1B and S1), we chose *CARDEL,* which had no prior cardiac link, to investigate its possible functional relevance in cardiac commitment (Figure 1C).

**Figure 1.**
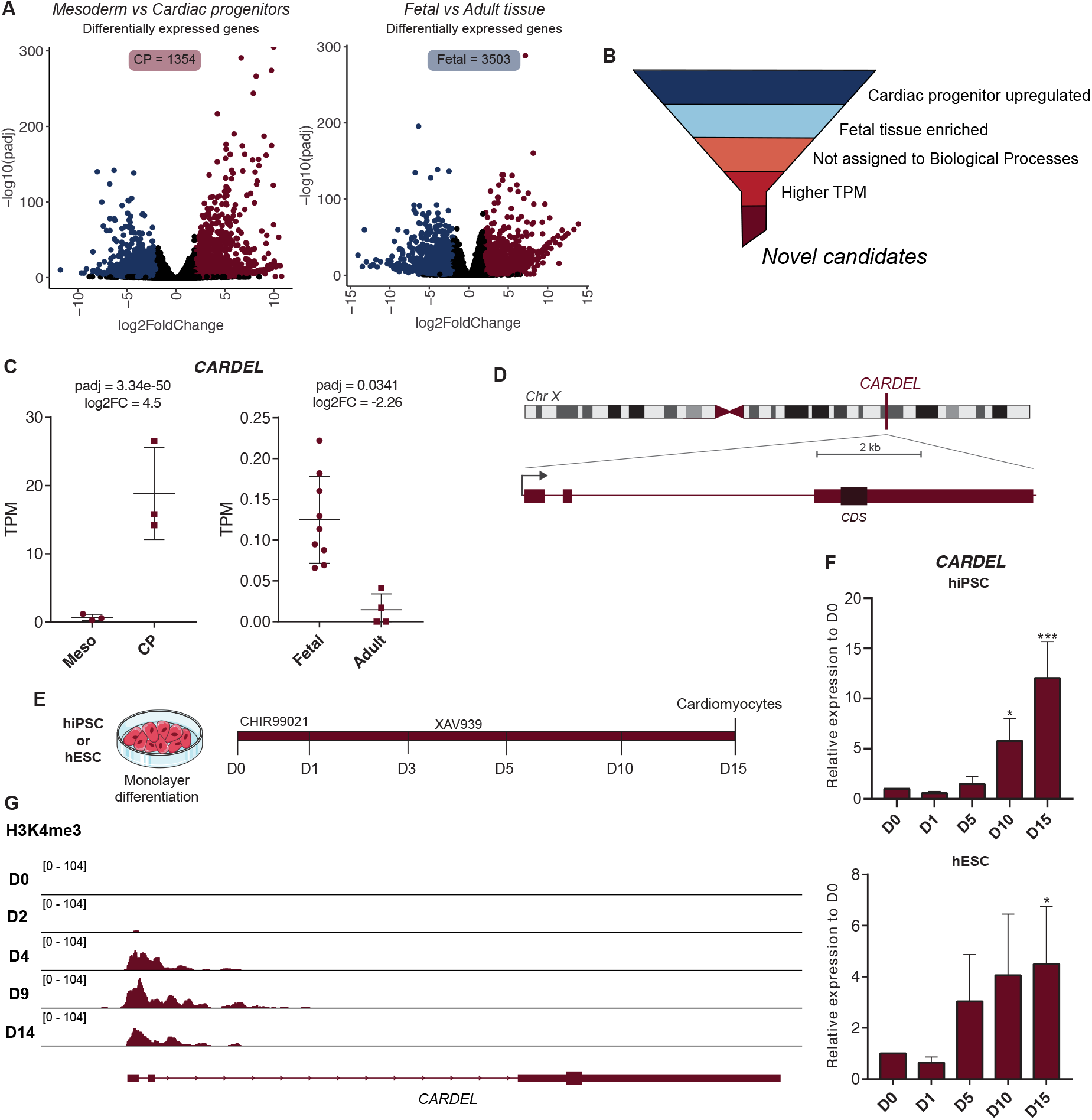
*CARDEL* is specifically expressed during the human cardiac specification. (A) RNA-seq analysis of mesoderm versus cardiac progenitor (CP) cells during cardiac IVD, and fetal versus adult heart tissues. Volcano plots show differentially expressed genes for each comparison: downregulated in blue, upregulated in red (padj < 0.05, −2 ≥ log2FC ≥ 2). (B) Schematic representation of the criteria used to find cardiac developmental genes candidates. (C) *CARDEL* expression in mesoderm and cardiac progenitor cells during cardiac IVD (left), and in fetal and adult heart tissues (right). TPM: transcripts per million. (D) Schematic representation of *CARDEL*’s gene locus at chromosome X showing its 3 exons and predicted coding sequence (CDS) position. Scale bar 2 kb. (E) Cardiac IVD protocol used to differentiate cardiomyocytes from hiPSC or hESC. The time of collection of differentiating cells for analysis is indicated in days. (F) *CARDEL* expression measured by RT-qPCR in differentiating cells from hiPSC or hESC. Mean with SD; One-way ANOVA followed by Dunnett’s multiple comparisons test. Each column compared with D0, *p < 0.05, ***p < 0.001. (G) ChIP-seq analysis of the active chromatin marker H3K4me3 (Paige et al., 2012) on *CARDEL* gene locus during cardiac IVD.

*CARDEL* (LINC00890; SERTM2) was named for (CAR)diac (DE)velopment (L)ong noncoding RNA. It is a transcript 4,612 nucleotides long that contains 3 exons and a polyadenylation sequence (Figure 1D). *CARDEL* is annotated in the latest Gencode version (GRCh38.p13) as a protein-coding gene, however, the coding potential of its short ORF still needs to be proven. Coding potential predicting algorithms, such as PhyloCSF and CPAT, classified *CARDEL*’s transcript as non-coding (Volders et al., 2019). On the other hand, the high conservation within its predicted coding sequence (CDS) is in favor of its annotation as proteincoding.

To further validate our observations with the RNA-seq data, we tested additional adult heart tissue samples for *CARDEL* expression by RT-qPCR and found undetectable expression in those samples (Figure S1). In corroboration with these results, we also interrogated a recently published database of over 900 RNA-seq samples (D’Antonio et al., 2022), which showed that *CARDEL* was enriched expressed in fetal-like cells when compared to adult heart tissue.

We therefore validated *CARDEL* expression in hPSC-derived cardiac lineage cells. We examined *CARDEL* temporal expression during cardiogenesis of both hESC and hiPSC, using one line of each (Figure 1E), and found that *CARDEL* expression coincides with cardiac lineage commitment, increasing from the cardiac mesoderm (D5) to cardiomyocytes (D15) (Figure 1F). Additional analysis of public ChIP-seq datasets (Paige et al., 2012) revealed an increase in the active chromatin marker H3K4me3 on *CARDEL* promoter during the IVD of hESC (Figure 1G), in agreement with the observed increase in expression in our data. Together, these results suggest that *CARDEL* is specifically expressed in early cardiac lineage cells. We therefore investigated the functional role of *CARDEL* during human cardiac specification.

### Cardiomyocyte differentiation and homeostasis were impaired in *CARDEL^KO^* cells

The functional activity of a genomic locus can be investigated at several levels: 1) DNA, such as regulatory sequences (e.g., promoters and enhancers); 2) RNA, as in non-coding roles either in cis or trans; or 3) protein. Considering that the *CARDEL* locus is uncharacterized to date, we first aimed to assess if *CARDEL* transcribed sequence was necessary for cardiac specification. To this end, we established two *CARDEL* knock-out (*CARDEL*^KO^) lines (H1 hESC and IPRN13.13 hiPSC) using CRISPR/Cas9 technology (Figure 2A). Cleavage in both gRNA target sequences could be detected by PCR using primers up and downstream to those sequences (KO-F/R), as well as internal primers (WT-F/R) (Figure 2A-B). *CARDEL*^KO^ lines showed normal karyotypes, typical pluripotent colonies, and expression of pluripotent genes, indicating that the lack of *CARDEL* expression does not affect pluripotency or genome stability (Figures 2C and S2). In addition, depletion of *CARDEL* expression was confirmed by RT-qPCR when *CARDEL*^KO^ cells were submitted to the cardiac IVD (Figure S2).

**Figure 2.**
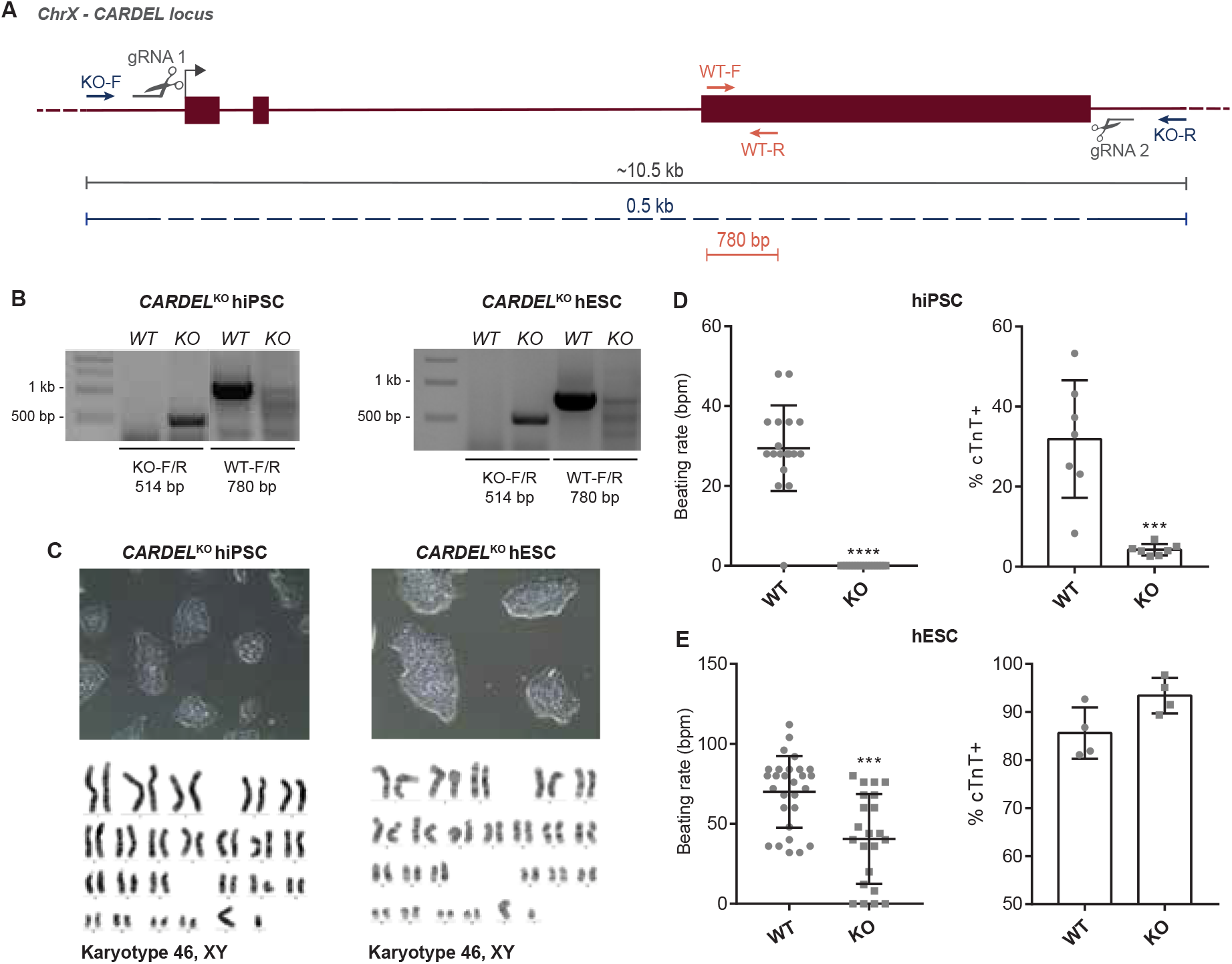
Cardiomyocyte differentiation and homeostasis were impaired in *CARDEL*^KO^ cells. (A) Schematic representation of the strategy used to knock-out *CARDEL* from hiPSC and hESC using CRISPR-Cas9. Positions of the gRNAs for Cas9 directed cleavage and of the oligos for confirmation PCR are indicated, as well as the expected fragment size. (B) PCR of genomic DNA confirming the knock-out on *CARDEL* locus in hiPSC and hESC. (C) Colony morphology of *CARDEL*^KO^ cells depicting pluripotency maintenance (top), and normal karyotype (bottom). (D, E) *CARDEL*^KO^ hiPSC- (D) and hESC-derived (E) cardiomyocytes beating rate (left) and quantification of cTnT positive cells by flow cytometry. Bpm: beats per minute. Mean with SD; Student’s unpaired t test analysis: ***p < 0.001, ****p < 0.0001.

Strikingly, *CARDEL*^KO^ hiPSC-derived cells did not beat as expected and showed very low cTnT positive cells when compared to wild-type (WT) hiPSC-derived cells (Figure 2D). Additionally, *CARDEL*^KO^ hESC-derived cells showed a decreased beating rate, but very similar cTnT positive cells (Figure 2E). It is worth noting that the baseline beating rate and cTnT positive yield in each hiPSC or hESC WT lines are different (Figure 2D-E), and the lower capacity of deriving cardiac cells from our parental hiPSC WT could explain the more striking deleterious phenotypes in these engineered cells. The lack or decrease of beating areas in the KO differentiating cells indicates that the absence of *CARDEL* expression impaired the cardiomyogenic differentiation of hPSC and suggests that *CARDEL* is essential for the success of cardiogenic commitment.

### Overexpression of *CARDEL* improves hPSC cardiomyogenic differentiation and alters cardiomyocyte functional properties

The hypothesis of *CARDEL* playing an essential role in cardiogenic specification led us to evaluate whether its controlled overexpression (OE) during differentiation would affect final cardiac lineage yield, especially in our parental hiPSC line that usually shows a poor cardiac yield. Thus, doxycycline-inducible pluripotent cell lines were established (i*CARDEL*^OE^ hESC and hiPSC) to express the *CARDEL* transcript containing its 3 exons (Figure 3A). The engineered cells showed normal karyotypes and *CARDEL* expression was confirmed by RT-qPCR after doxycycline induction (Figure S3). We tested different windows of induction and measured cTnT positive cells as an output quantification. Interestingly, final cardiomyocyte yield was negatively affected in many tested conditions but one. One possible explanation could be that *CARDEL* expression has to be carefully controlled during cellular differentiation, and exogenous *CARDEL* placed at the wrong time negatively affects the cardiogenic commitment. i*CARDEL*^OE^ hiPSCs submitted to the cardiomyogenic IVD and induced expression from day 5 to day 10 (D5-10) showed increased number of cTnT positive cells and a larger fraction of beating area when compared to non-induced cells (Figure 3C). Curiously, endogenous *CARDEL* expression starts increasing at the cardiac mesoderm stage at day 5 (Figure 1F), and the induction from D5-10 mimics its physiological timeline (Figure 3B). These results clearly indicate an improvement in the cardiac differentiation efficiency as an effect of *CARDEL* overexpression. On the other hand, i*CARDEL*^OE^ hESC-derived cardiomyocytes did not show significant differences in beating areas or cTnT positive cells when induced with doxycycline. This might have happened because is more difficult to see improvement over the high cardiomyocyte yield (up to 80%) that noninduced hESCs usually have (Figure 3D). Moreover, both i*CARDEL*^OE^ hiPSC- and hESC-derived cardiomyocytes showed faster beating rates when compared to the non-induced control (Figure 3E-F, Video S1-2). Together, these results strongly suggest that overexpression of *CARDEL* during cardiomyocyte specification can improve the final differentiation output and alter functional properties of those cells.

**Figure 3.**
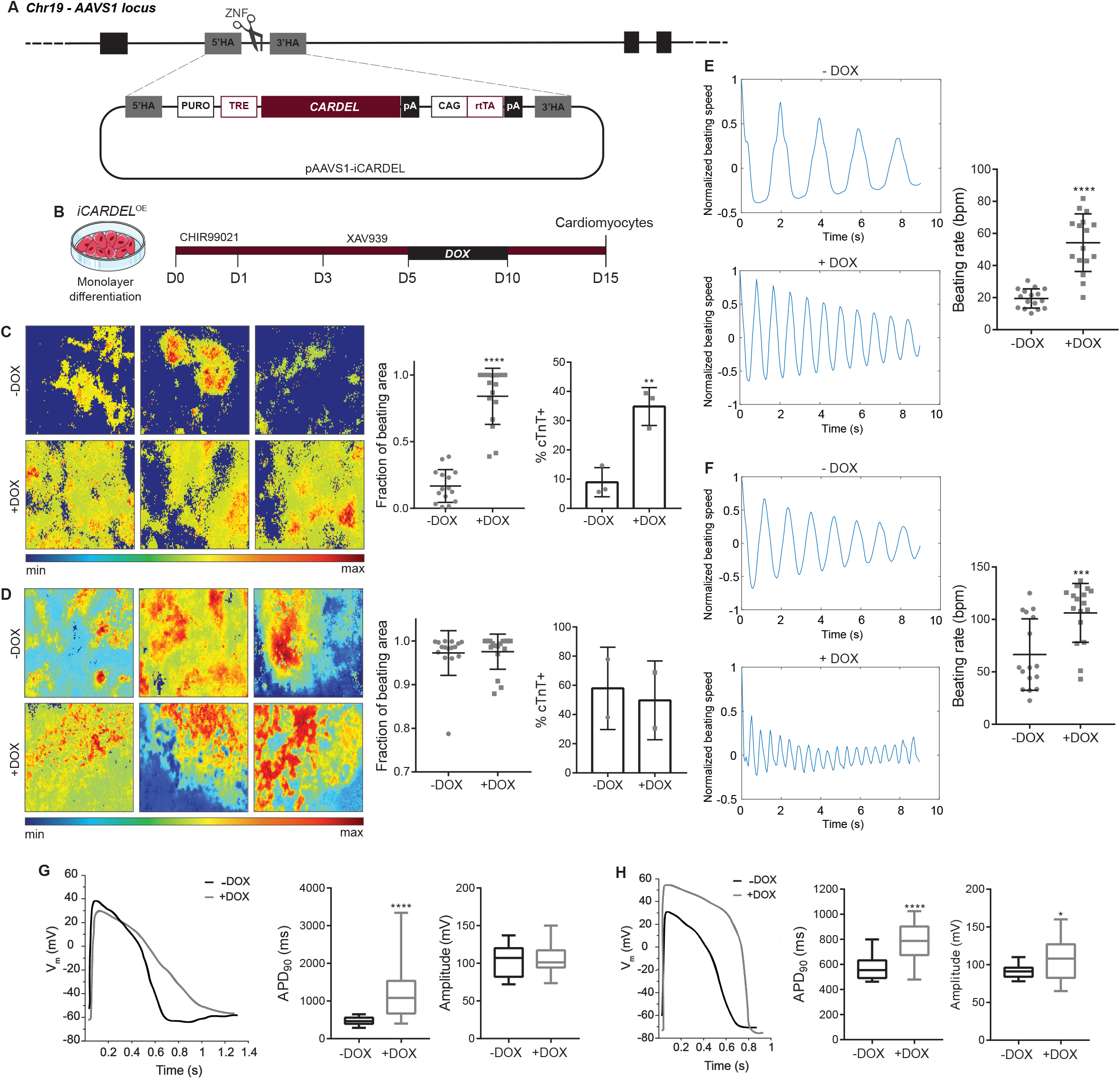
*CARDEL* overexpression improves hPSC cardiomyogenic differentiation and alters cardiomyocyte functional properties. (A) Schematic representation of dox-inducible system knock-in on AAVS1 locus. (B) Cardiac IVD protocol used to differentiate cardiomyocytes from dox-inducible i*CARDEL*^OE^ cells. The time is indicated in days. (C, D) i*CARDEL*^OE^ hiPSC- (C) and hESC-derived (D) cardiomyocytes differentiated in control (−DOX) or induced (+DOX) conditions. Representative images of the intensity of beating areas according to the heatmap scale in the bottom (left). Quantification of beating areas (center). Quantification of cTnT positive cells by flow cytometry (right). (E, F) Representative plots of beating frequency (left) and quantification of beating rate in i*CARDEL*^OE^ hiPSC- (E) and hESC-derived (F) cardiomyocytes differentiated in control (−DOX) or induced (+DOX) conditions. (G, H) Action potential profiles in *iCARDEL*^OE^ (G) hiPSC- and (H) hESC-derived ventricular cardiomyocytes in control (−DOX) or induced (+DOX) conditions. Mean with SD; Student’s unpaired t test analysis: * p < 0.05, **p < 0.01, ***p < 0.001, ****p < 0.0001.

### *CARDEL* expression contributes to physiological and molecular changes during cardiomyocyte differentiation

Since we observed an impressive increase in the beating rate of i*CARDEL*^OE^ cardiomyocytes, we further explored their functional properties. Whole cell patch clamp assays in i*CARDEL*^OE^ hPSC-derived cardiomyocytes were performed. After induction with doxycycline from D5-10, both i*CARDEL*^OE^ hiPSC- and hESC-derived cardiomyocytes showed altered action potential (AP) profiles and overall longer action potential duration (APD) in induced cells. The APD at 90% of repolarization (APD_90_) of the ventricular type of cells was 476.6 ± 22.8 ms in −DOX and 1169.0 ± 129.4 ms in +DOX for i*CARDEL*^OE^ hiPSC, and 571.0 ± 21.5 ms in −DOX and 775.3 ± 36.9 ms in +DOX for *iCARDEL^OE^* hESC (Figure 3G-H). Regarding each cardiomyocyte subtype, i*CARDEL*^OE^ hiPSC-derived atrial and nodal cells also showed increased APD_90_; and ventricular and nodal cells showed decreased maximum upstroke velocity (dV/dt_max_), suggesting altered contractility properties. i*CARDEL*^OE^ hESC-derived ventricular and atrial cells showed alterations in the AP amplitude (Figure S3).

To better understand the phenotypic alterations observed after induction of *CARDEL* during differentiation, we performed RNA-seq of polysome-bound RNAs from i*CARDEL*^OE^ cardiac progenitors at day 10 and cardiomyocytes at day 15 under control and induced conditions. Compared to non-induced control cells, 195 genes were upregulated and 516 downregulated on day 10. On day 15, 393 genes were upregulated and 522 were downregulated (Figure 4A and Table S3). Accounted together, a larger number of differentially expressed genes were downregulated in induced (+DOX) conditions. However, Genome Ontology (GO) analysis of these sets of genes showed more general terms related to morphogenesis and development. Interestingly, upregulated genes in either day 10 and 15 were related to many cardiac specific GO terms, such as “regulation of heart contraction”, “cardiac conduction”, “cardiac muscle tissue development” and “heart development” (Figure 4B). Some genes assigned in these terms are depicted in heatmaps for comparison of expression level between the four conditions analyzed (Figure 4C-D). Cluster analysis shows that the expression levels of many genes upregulated in day 15 +DOX are more similar to day 10 +DOX than to non-induced cells in day 15 (Figure 4C). This suggests that *CARDEL* overexpression modulates the cardiac molecular profile strongly enough to differ either early (day 10) and late (day 15) cardiomyocytes from those non-induced. Among the upregulated genes, critical cardiac development transcription factors, such as *NXK2-5* and *IRX3* can be found. Also, an extensive list of myosins and sarcomere structural proteins appeared as upregulated in +DOX cardiomyocytes, including *MYH6, MYH7, MYL3, TNNT2* and *TNNC1.* Genes related to cardiac signaling and homeostasis were also upregulated in these cells, as the hormone atrial natriuretic peptide (*NPPA*) and the protease corin (*CORIN*), which converts the pro-atrial natriuretic peptide to its biologically active form (Figure 4D).

**Figure 4.**
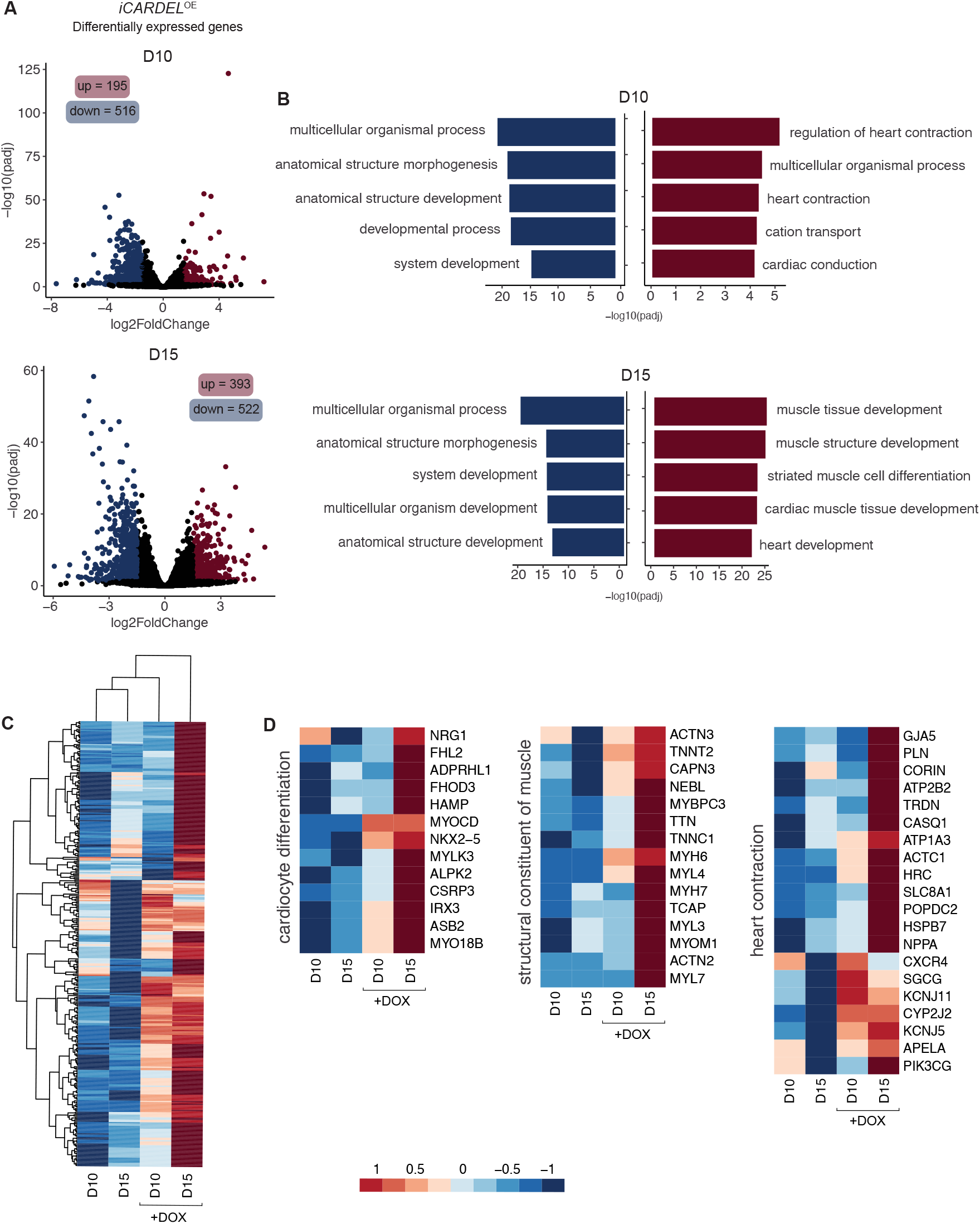
*CARDEL* contributes to molecular changes during cardiomyocyte differentiation. (A) RNA-seq analysis of i*CARDEL*^OE^ hiPSC-derived cardiomyocytes differentiated in control (−DOX) or induced (+DOX) conditions. Volcano plots show differentially expressed genes for −DOX versus +DOX: upregulated in red, downregulated in blue (padj < 0.05, −1.5 ≥ log2FC ≥ 1.5). (B) Genome Ontology (GO) analysis of upregulated (red) and downregulated (blue) sets of genes. Five most representative GO terms are shown. (C) Heatmap and cluster of upregulated genes in D15. (D) Heatmap of upregulated genes in D15 related to specific GO terms as indicated. Values are plotted as log2 of normalized counts by DESeq2.

These results provide robust evidence that *CARDEL* overexpression during cardiac lineage commitment alters cardiomyogenic differentiation at both physiological and molecular levels, contributing to the distinct phenotype observed and suggesting a role for this gene in heart contraction homeostasis during development.

## Discussion

In this report, we showed that *CARDEL* expression is tightly controlled in differentiating cells and plays a crucial role in the physiology of derived cardiomyocytes.

*In vitro* cardiac differentiation of stem cells is an important tool for the study of heart developmental mechanisms (Hartman et al., 2016). Studies using hPSC have led the way to understanding how gene expression is controlled during human development as the access to fetal-derived biological material is very limited. Analysis of mammalian stem cell transcriptomes during *in vitro* cardiomyocyte differentiation revealed thousands of differentially expressed genes, which are tightly and timely controlled (Fu et al., 2018; den Hartogh et al., 2016; Kurian et al., 2015; Li et al., 2015; Pereira et al., 2019; Tompkins et al., 2016; Xu et al., 2009). However, while a huge set of biological data has been generated by high throughput approaches, the lack of functional characterization hampers reliable identification of genes correlated with specific biological processes. Considering that functional annotation is critical to integrate and analyze complex biological data (Uchida et al., 2009), the characterization of critical developmental players contributes to countless fields, such as the comprehension of heart congenital diseases and the application of cellular and genetic therapies in regenerative approaches.

Here, our main goal was to find and functionally characterize a cardiac developmental gene. Combining a temporal evaluation of the human cardiac specification and tissue expression, we develop a strategy to search for genes exclusively expressed during differentiation. Our stringent approach found *CARDEL* as one very interesting candidate based on its robust expression in independent and distinct *in vitro* differentiation protocols and cell lines (Figure 1). As an uncharacterized gene, we chose a wider genetic approach to begin with its functional characterization. The deletion of its entire locus, including all exons and introns but excluding its promoter region, showed that *CARDEL* is required for a good cardiac output after differentiation (Figure 2). The importance of noncoding DNA elements (e.g., enhancers and insulators) in a specific locus is not clearly discriminated from the importance of the RNA transcript itself or the act of transcription in a deletion-based knockout strategy (Kopp and Mendell, 2018; Nakagawa, 2016). For this reason, we performed an unbiased and independent analysis of the biological consequence of *CARDEL* transcript expression. Using a drug inducible system to temporally control *CARDEL* expression, we showed that the *CARDEL* transcript comprising its 3 exons, when expressed from a completely independent locus, was able to promote cardiomyocyte differentiation and phenotype (Figures 3 and 4).

*CARDEL* annotation shows high inter-species conservation in a predicted coding sequence (CDS), which classifies it as a protein-coding gene. However, its coding potential still needs to be proved. The stringent and arbitrary criteria traditionally used for prediction of protein-coding open reading frames (ORFs), e.g., 300 nucleotides or 100 amino acids as a size cutoff and AUG start codon, have led to misannotation of many RNAs that contain small ORFs and potentially encode micropeptides (Hartford and Lal, 2020; Yeasmin et al., 2018). Also, detection sensitivity and limited database for spectra search in mass spectrometry approaches have always represented challenges for the identification of small peptides. Recent and emerging advancements in bioinformatics, proteomics and transcriptomics have made the identification of new potential small ORFs possible, including in the heart (Abascal et al., 2020; van Heesch et al., 2019; Housman and Ulitsky, 2016; Mackowiak et al., 2015; Wang et al., 2017). To date, no peptide derived from *CARDEL* locus has been identified and biologically characterized. Additionally, the presence of an ORF does not necessarily mean that the transcript is being translated. For instance, the LINC00261 was found as dynamically regulated during pancreatic differentiation of stem cells, being predominantly cytosolic and associated with ribosomes based on ribosome profiling data (Gaertner et al., 2020). However, disruption of each of LINC00261’s seven identified ORFs excluded the requirement of the derived peptides and suggested that it is the RNA that is involved in the differentiation. Myosin heavy chain 7b (MYH7b) is also an interesting example of a locus encoding several potential active molecules: the protein myosin, a miRNA and a lncRNA. However, in the heart, the locus does not produce a protein due to an exon-skipping mechanism. It was recently shown that it is the lncRNA that has a regulatory role affecting the transcriptional landscape of human cardiomyocytes, independent of the miRNA (Broadwell et al., 2021).

Many studies have found lncRNAs as critical players during the cardiac development (Devaux et al., 2015; Kurian et al., 2015; Li et al., 2017; Lopez-Pajares, 2016; Pereira et al., 2020). Braveheart, Fendrr and Carmn are examples of lncRNA already functionally described, which loss of expression was associated with a significant impairment of heart development (Grote et al., 2013; Klattenhoff et al., 2013; Ounzain et al., 2015). Therefore, *CARDEL’s* RNA could be acting in a non-coding mechanism. Nevertheless, *CARDEL* gene product is shown here as developmentally expressed and required for cardiac specification, which likely acts in trans. *CARDEL* overexpression in hiPSC showed a remarkable improvement in the final cardiac output (Fig 3). Additionally, the most consistent observed phenotype was the increase in the beating rate, which occurred in both cell lines studied. *CARDEL* overexpressing cells also showed increased APD90 and upregulation of many heart contraction genes, such as cardiac ion channels and sarcomere structural genes. The development of mature sarcomeres and sarcoplasmic reticulum allow faster and efficient contraction, leading to better electric coupling of cardiomyocytes (Buijtendijk et al., 2020). Therefore, these results suggest that, beyond improving differentiation, *CARDEL* is also acting in cardiomyocyte physiological and functional properties.

Cardiac hypertrophy is a pathological process that is accompanied by molecular changes that resemble those observed during cardiac development (Dirkx et al., 2013). In addition, arrhythmia is usually associated with cardiac hypertrophy and other cardiac conditions. It is important to note that *CARDEL* was found enriched in fetal tissue and cardiac developmental staged cells (Figure 1), as well as its overexpression caused an increase in the cardiomyocyte beating rates (Figure 3). A systematic study using RNA-seq data from 28 hypertrophic cardiomyopathy patients and 9 healthy donors identified *CARDEL* as upregulated in diseased tissues (Gao et al., 2020). On the other hand, other studies comparing human left ventricles of dilated or ischemic cardiomyopathy patients did not find *CARDEL* among the differentially expressed genes. So, it is possible that *CARDEL* expression can be related to certain cardiac diseases, but this association remains inconclusive.

While further research is still needed to clarify the exact molecular mechanisms underlying our observations, this study adds a new piece to the intricate gene network regulated during the cardiac commitment. Ultimately, this effort enhances our knowledge about human development and helps to overcome the remaining challenges to understand and treat heart diseases.

## Experimental procedures

### Resource availability

#### Corresponding author

Further information and requests for resources and reagents should be directed to and will be fulfilled by the lead contact, Dr. Isabela T. Pereira.

### Materials availability

Plasmids generated in this study are available upon request.

### Data and code availability

RNA-Seq data generated in this study have been deposited in the NCBI Sequence Read Archive under accession number PRJNA926378 and are publicly available as of the date of publication.

This paper does not report original code.

Any additional information required to reanalyze the data reported in this paper is available from the lead contact upon request.

### Method details

#### Cell culture and cardiomyocyte differentiation

H1 hESC line and IPRN13.13 iPSC line are male and present normal karyotypes. Cells were cultured on Matrigel-coated dishes using either mTeSR-1 or TeSR-E8 medium (Stem Cell Technologies). The medium was replaced daily until cells get 70-80% confluence. Cells were maintained in a 37°C, 5% CO_2_ incubator.

Both cell lines were submitted to a protocol based on (Lian et al., 2012), with a few modifications. Briefly, 5 × 10^5^ cells/well were seeded in Matrigel-coated 12-well plate in mTeSR-1 + 10 uM Y27632; this time point corresponds to day - 3 of differentiation. The medium was replaced with 2 ml of mTeSR-1 per well on days −2 and −1. On day 0, 2 ml of RPMI/B-27 minus insulin medium supplemented with 12 μM CHIR99021 were added in each well. After 24 h (day 1), the medium was replaced with 2 ml of RPMI/B-27 minus insulin. On day 3 of differentiation, 1 ml of conditioned medium from the well was mixed with 1 ml of fresh RPMI/B-27 minus insulin medium and supplemented with 10 uM XAV939 to be replaced in the well. On day 5, the medium was replaced with 2 ml of fresh RPMI/B-27 minus insulin; and on day 7, day 10 and day 13 the medium was replaced with 2 ml of RPMI/B-27. When maintained until day 30 of differentiation, the medium was replaced with RPMI/B-27 every 3-4 days. Beating areas were expected around day 10. In the doxycycline-inducible lines, 1 ug/ml doxycycline was added on day 5 and day 7 to complete a day 5 to 10 induction.

Beating areas were recorded using AxioVs40 V4.8.2.0 on an inverted microscope (Carl Zeiss MicroImaging), from two distinct areas within the same well, three wells per differentiation replicate, three (hiPSC) or two (hESC) differentiation replicates in total. Beating area and rate were quantified using Motion Tracking software (Huebsch et al., 2015).

#### Doxycycline-inducible expressing and knock-out lines

For the doxycycline-inducible expressing lines, *CARDEL* sequence (4.5 kb) was amplified from the DKFZp686D0853 clone (gift from Dr. S. Weimann, German Cancer Research Center, Germany) and cloned into the pDONOR221 vector, and then into the AAVS1-TRE-GW-rtTA vector through Gateway methodology. hPSCs were transfected using 600 ng of AAVS1-TRE-*GW-CARDEL-rtTA* construct, 200 ng of ZFN(R) and ZFN(L), for the insertion of the doxycycline-inducible system into the AAVS1 *locus.* Positive transfected cells were selected with 0.2 μg/ml puromycin for 7 days and single-colony cloned. Doxycycline-inducible expression was confirmed by RT-qPCR.

For the knock-out lines, two guide RNAs were designed neighboring the *CARDEL locus* and were cloned into the pSpCas9(BB)-2A-RFP vector. hPSCs were transfected using 500 ng of each construct. Mutated colonies were screened by PCR after single-colony cloning. Genotyping was performed using genomic DNA and primers for the *CARDEL* internal sequence (WT-F: GCTGGGCAGGAACCTTACAA, WT-R: TGTTTGAGCCGAAGGAGCAT; detection of 780 bp wild-type allele) and for the *CARDEL* neighboring sequences (KO-F: GGGCCTTTGACTACAAATGGAT, KO-R: GCAGAGGAATGTGGAAGGCT; detection of 514 bp deleted allele and 10.500 bp wild-type allele).

#### Karyotype

The new derived cell lines were seeded in 6-well plates and incubated with 0.32 ug/ml colchicine for 90 min when reached 90% confluence. After washing with 1X PBS, cells were incubated with 1X PBS for 10 min at 37 °C and then detached using a scraper. Then, cells were collected and centrifuged at 800 xg for 5 min. The cell pellet was resuspended in 57 mM KCl for 10 min incubation at 37°C. Cells were then fixed using a solution containing 75% methanol and 25% acetic acid for 10 min at −20°C, followed by two rounds of washes in a solution of 66,6% methanol and 33,3% acetic acid. Fixed cells were used to prepare slides and karyotype analysis.

#### Flow cytometry

hPSC-derived cardiomyocytes were dissociated using trypsin-EDTA (0.05%) and fixed using 4% paraformaldehyde and 90% methanol. Fixed cells were incubated with 0.5% PBS/BSA, 0.1% Triton-X and 1:100 primary antibodies for cTnT (cardiac isoform Ab-1 mouse, Thermo Scientific™, cat. #MS-295-P0). After washing, cells were incubated with 0.5% PBS/BSA, 0.1% Triton-X and 1:1000 secondary antibodies. Analyses were carried out using a FACSCanto II flow cytometer and FlowJo software.

#### RNA isolation and RT-qPCR

Total RNA of cells in distinct time-points of differentiation was isolated using TRIreagent (Sigma-Aldrich) according to manufacturer’s instructions. cDNA synthesis was performed using ImProm-II™ Reverse Transcriptional System (A3800 - Promega) with 1 ug of RNA and RT-qPCR reactions were performed using GoTaq® qPCR Master Mix (A6002 - Promega). Analyses were carried out in the QuantStudio™ 5 Real-Time PCR System. GAPDH or POLR2A expression was used as an endogenous control.

#### Whole cell patch clamp

Action potentials from day 30 cardiomyocytes were measured using the whole-cell patch clamp technique in current clamp mode at room temperature. Pipettes (2-3 MΩ) were filled with a pipette solution containing (in mM): NaCl 5, KCl 150, CaCl2 2, HEPES 10, EGTA 5, MgATP 5 (adjusted to pH 7.2 with KOH). The bath solution will consist of (in mM): NaCl 150, KCl 5.4, CaCl2 1.8, MgCl2 1, HEPES 15, glucose 15 (adjusted to pH 7.4 with NaOH). All electrophysiological measurements were carried out with an Axopatch 200B amplifier and Axon Digitata 1320A A/D converter driven by a pCLAMP system (Digidata A/D and D/A boards and pCLAMP 9.2, Molecular Devices, Sunnyvale, CA). Data were analyzed with Clampfit. All records were filtered with cutoff frequencies designed to avoid aliasing and were digitized at speeds at least five times the filter cutoff frequency, generally 2 kHz and 10 kHz, respectively. The APs were recorded under pulses of 3 ms in duration, 1.2-fold the threshold intensity at a stimulation rate of 0.5 Hz. Analog and P/4 methods were used for leak and capacity transient cancellation. Series resistance was partially compensated by feedback circuitry.

#### RNA-seq and data analysis

RNA-seq data of *in vitro* cardiac differentiation was previously generated by our group (Pereira et al., 2018) and is available at NCBI Sequence Read Archive, accession number SRP150416. RNA-seq data heart tissues were acquired from European Nucleotide Archive (ENA) (http://www.ebi.ac.uk/ena). Accession numbers for adult heart tissue data: ERR315356, ERR315430, ERR315367 and ERR315331. Project number for fetal heart tissue data: PRJEB27811 (Pervolaraki et al., 2018). Quality control, read mapping, and quantification was performed using nf-core/rnaseq (v. 3.8.1) pipeline. Reads were mapped to hg38/GRCh38 and quantified with Salmon using the Gencode v38 annotation. Significantly differentially expressed genes were calculated using DESeq2 (v.1.24.0) (Love et al., 2014) with an adjusted p-value cutoff of 0.05 and log2FoldChange cutoff of 2.

RNA-seq data of *CARDEL* overexpression was generated using i*CARDEL*^OE^ hiPSC cell line in control (−DOX) and induced (+DOX, 1 ug/ml doxycycline was added on day 5 and day 7 to complete a day 5 to 10 induction) conditions. Quality assessment and trimming of reads was done with FastQC and Trim Galore (v.0.4.0) (Krueger). Reads were mapped to hg38/GRCh38 with HISAT2 (v.2.1.0) (Kim et al., 2015) and counted with HTSeq (v.0.11.1) (Anders et al., 2015). Significantly differentially expressed genes were calculated with DESeq2 (v.1.24.0) (Love et al., 2014) with an adjusted p-value cutoff of 0.05 and log2FoldChange cutoff of 1.5. Normalization of reads by RPKM was performed with edgeR (v.3.40.0) (Robinson et al., 2010).

ChIP-seq data is described in Paige et al., 2012 and is available at Gene Expression Omnibus under accession number GSE35583. BigWig files were retrieved from ENCODE (ENCODE4 v1.5.1 GRCh38 processed data).

Gene Ontology (GO) analysis was performed using gProfiler (Raudvere et al., 2019).

#### Data and statistical analysis

Graphed data are expressed as mean ± SD. Statistical analysis was performed with Prism 8.0 software (GraphPad Software). For comparison between multiple groups, a one-way ANOVA followed by Dunnett’s multiple comparison test was performed as appropriate. Comparisons between two mean values were performed with a Student’s unpaired t-test. A p-value of less than 0.05 was considered statistically significant. For RNA-seq data, DESeq2 (v.1.24.0) (Love et al., 2014) was used for statistical analysis with an adjusted p-value less than 0.05 considered significant. Patch clamp data recording was performed by a collaborator who was blinded to the experimental conditions.

## Supporting information

DataS1

DataS2

DataS3

## Acknowledgements

We would like to thank Dr Paul J. Gadue from University of Pennsylvania, USA, for kindly sharing the ZFN expression vectors and AAVS1 targeting construct. We also would like to thank all the staff of the Carlos Chagas Institute (FIOCRUZ-PR) for the laboratory and administrative support. This work was supported by CNPq: National Center of Science and Technology for Regenerative Medicine/REGENERA grant 465656/2014-5, CNPq PROEP/ICC grant 442353/2019-7, CAPES grant INCT-REGENERA 88887.136364/2017-00, and NIH grants 1R56HL162208 and R01-HL134791. I.T.P., R.G. and A.H. received fellowships from CAPES and B.D. from CNPq.

## Author contributions

I.T.P., S.S.K.S., M.K. and B.D. developed concept and designed experiments. I.T.P., R.G., M.L. and H.A.N.S. performed the experiments. I.T.P. and A.H. performed the bioinformatic analyses. S.C.D. coordinated patch-clamp experiments. M.K. and B.D. supervised all aspects of the project. I.T.P. wrote the manuscript. All authors read and approved the final manuscript.

---

Data S1. Differential expression analysis of (A) mesoderm versus cardiac progenitor cells in the IVD of hPSCs, and (B) fetal versus adult heart tissues. Results from DESeq2 analysis of RNA-seq data. Each comparison table contains gene_id, gene_name, log2FoldChange between sample groups and adjusted p values. (C) List of candidate genes. The table contains gene_id and gene_name of genes in the intersection between enriched in cardiac progenitors and enriched in fetal heart tissues. Related to Figure 1.

Data S2. (A) Gene Ontology analysis of list of candidates from Data S1-C. The table contains the output from gProfiler GO analysis with “term_name” for GO category and “intersections” for genes present in the dataset. (B) Filtered candidate genes list. The table contains gene_id and gene_name of genes from Data S1-C not assigned to GO categories. Related to Figure 1.

Data S3. Differential expression analysis of i*CARDEL*^OE^ hiPSC-derived cardiomyocytes differentiated in control (−DOX) versus induced (+DOX) conditions at (A) D10 and (B) D15. Results from DESeq2 analysis of RNA-seq data. The table contains gene_id, gene_name, log2FoldChange between sample groups, adjusted p values and normalized counts. Related to Figure 4.

Video S1. i*CARDEL*^OE^ hiPSC-derived cardiomyocytes differentiated in control (−DOX) conditions at D15.

Video S2. i*CARDEL*^OE^ hiPSC-derived cardiomyocytes differentiated in induced (+DOX) conditions at D15.

