## Supplemental Figures for "Cardiac Development Long non-coding RNA (CARDEL) is activated during human heart development and contributes to cardiac specification and homeostasis"

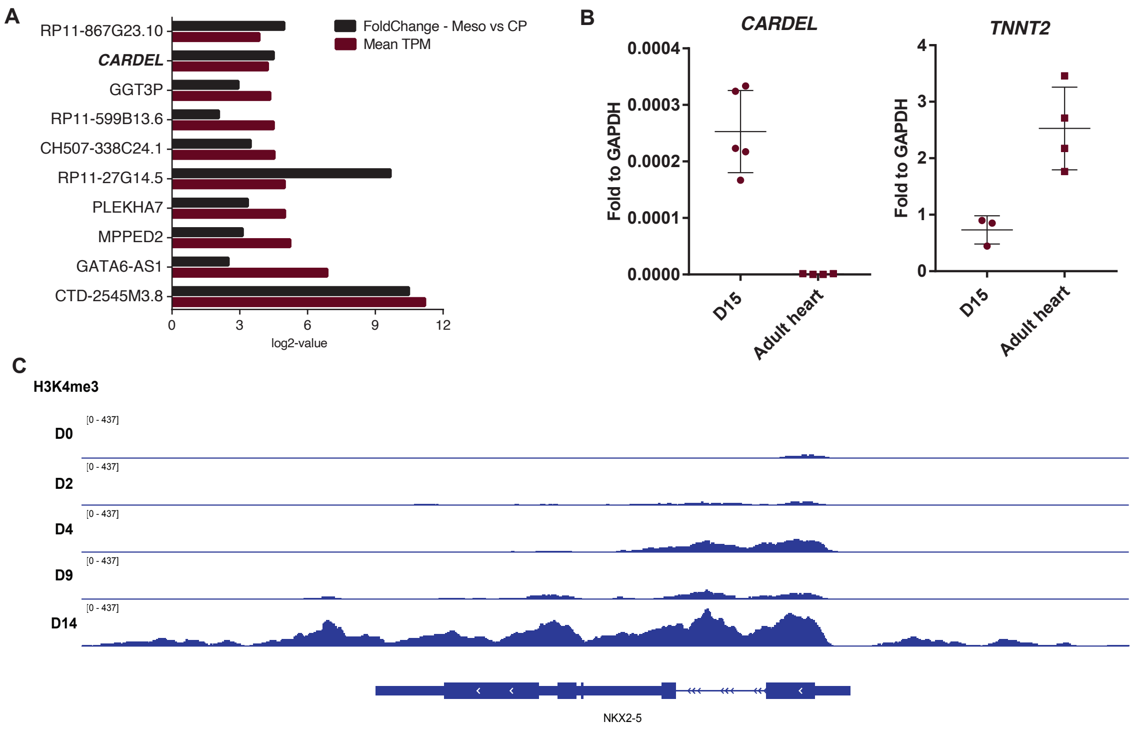


Figure S1. (A) Top 10 genes with higher TPM (transcripts per million) among those upregulated in CP cells when compared to mesoderm cells. (B) CARDEL expression measured by RT-qPCR in additional adult heart samples. CARDEL expression in hESC-derived cardiomyocytes (D15) and TNNT2 expression are shown as a reference. (C) ChIP-seq analysis of the active chromatin marker H3K4me3 (Paige et al., 2012) on NKX2-5 gene locus is shown as an example of a cardiac gene being activated during in vitro cardiac differentiation. Related to Figure 1.

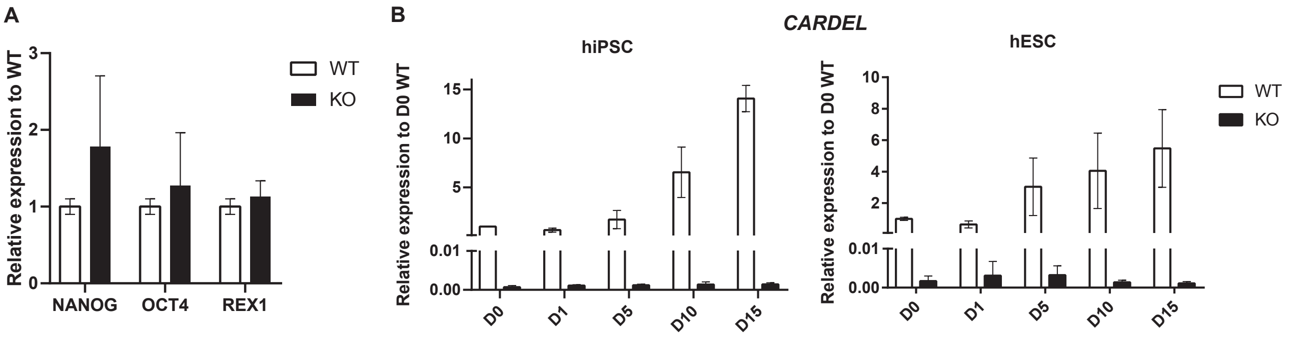


Figure S2. (A) Expression of pluripotency gene markers measured by RT-qPCR in wild-type (WT) and CARDELKO hiPSCs. (B) CARDEL expression measured by RT-qPCR in wild-type (WT, same as in Figure 1F) and CARDELKO cells during cardiac differentiation in hiPSC and hESC. Related to Figure 2.

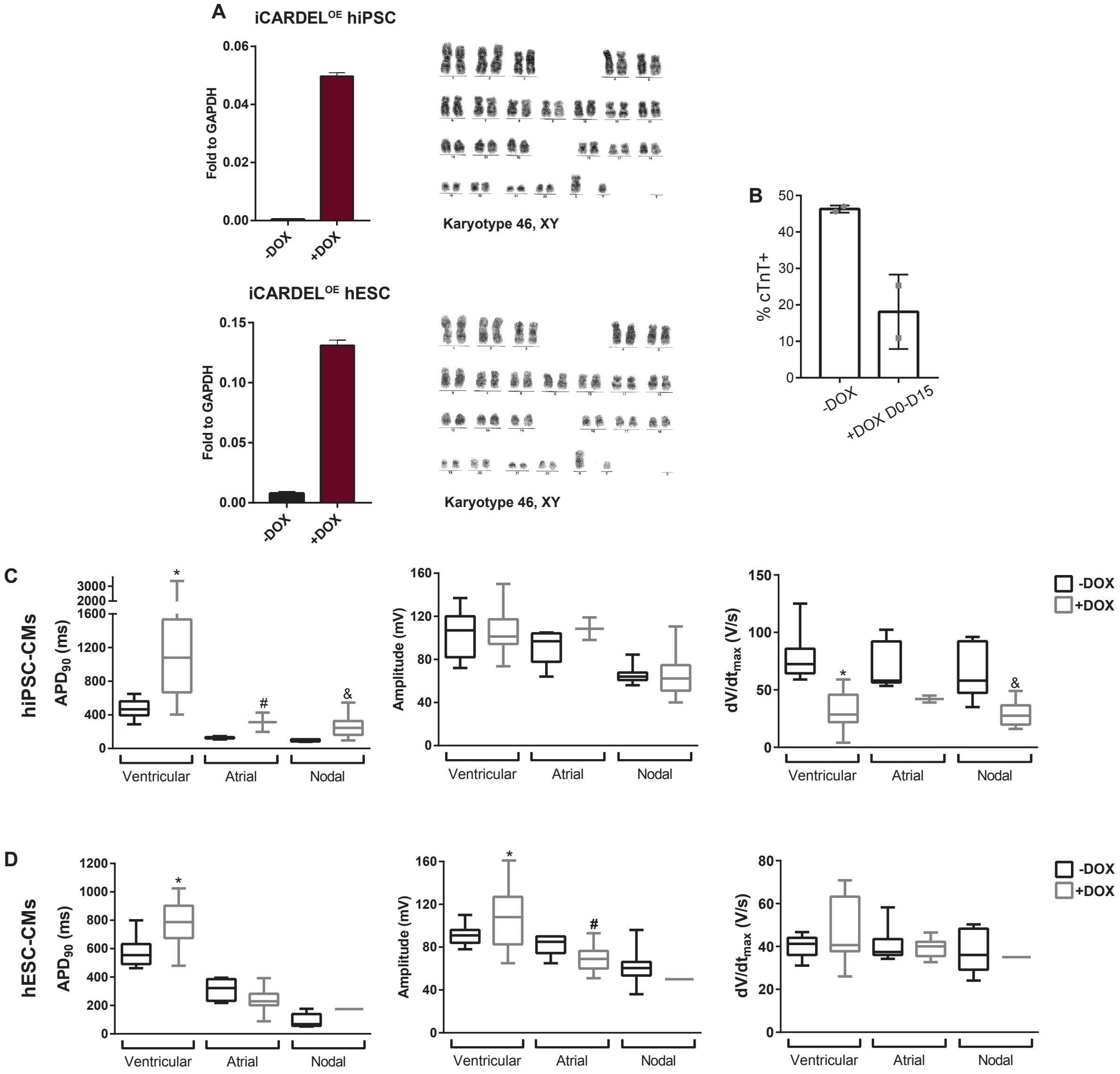


Figure S3. (A) CARDEL expression measured by RT-qPCR before and after dox induction on selected iCARDELOE clones (left), and their normal karyotypes (right). (B) Quantification of cTnT positive cells by flow cytometry after cardiac differentiation in control (-DOX) or induction (+DOX) from day 0 to day 15. (C) Quantification of cTnT positive cells by flow cytometry after cardiac differentiation in control (-DOX) or induction (+DOX) in hESC WT. (D, E) Quantification of electrophysiological properties in ventricular, atrial and nodal iCARDELOE (D) hiPSC- and (E) hESC-derived cardiomyocytes in control (-DOX) or induced (+DOX) conditions. Student's unpaired t test analysis: *ventricular, #atrial, &nodal for $p < 0.05$ when compared to -DOX. Related to Figure 3.
